# In bacterial membranes lipid II changes the stability of pores formed by the antimicrobial peptide nisin

**DOI:** 10.1101/2024.02.14.580365

**Authors:** Miranda S. Sheridan, Preeti Pandey, Ulrich H. E. Hansmann

## Abstract

Resistance to available antibiotics poses a growing challenge to modern medicine as this often disallows infections to be controlled. This problem can only be alleviated by developing new drugs. Nisin, a natural lantibiotic with broad antimicrobial activity, has shown promise as a potential candidate for combating antibiotic-resistant bacteria. However, nisin is poorly soluble and barely stable at physiological pH, which, despite attempts to address these issues through mutant design, has restricted its use as a antibacterial drug. Therefore, gaining a deeper understanding of the antimicrobial effectiveness, which relies in part on its ability to form pores, is crucial for finding innovative ways to manage infections caused by resistant bacteria. Using large-scale molecular dynamics simulations we find that the bacterial membrane specific lipid II increases the stability of pores formed by nisin, and that the interplay of nisin and lipid II reduces the overall integrity of bacterial membranes by changing local thickness and viscosity.

## I. INTRODUCTION

The resistance to available antibiotics has become an growing problem in healthcare, significantly limiting the range of drugs available for controlling bacterial infections.^1^ An intriguing candidate in the urgent quest for new types of antibiotics is the antimicrobial peptide nisin, which is synthesized by the gram-positive bacterium *Lactococcus lactis*. Nisin has a wide range of antibacterial effects and kills gram-positive bacteria even at low concentrations.^2^ This lantibiotic is a 34-residue long positively charged peptide containing uncommon amino acids such as dehydroalanine (DHA) and dehydrobutyrine (DHB), D-alanine (DAL), and four aminobutyric acids (ABA), forming five thioether rings in the peptide, as illustrated in **Figure 1a**. Nisin eliminates bacteria by employing two different modes of action. In gram-positive bacteria, the precursors UDP-N-acetyl-glucosamine and N-acetylmuramic-acid-pentapeptide coalesce with a prenyl chain and two phosphate atoms to form lipid II, see **Figure 1b**. In this form, lipid II transports the precursors from the cytoplasm to the exterior, where they are added to the peptidoglycan layer, enabling the construction of bacterial cell walls.^3^ By binding of the first twelve residues of nisin to the pyrophosphate moiety of lipid II (drawn in red and white in **Figure 1b**) nisin interferes with the biosynthesis of the cell wall.^4^ On the other hand, lipid II acts after binding as an anchor, allowing nisin to move into the membrane, where eight nisin chains may associate into a pore bound to four lipid II chains (**Figure 1c**). The presence of such openings (pores) compromises the integrity of the membrane bilayer, which acts as a barrier separating the interior of the cell from its surroundings.^5^ It is this double mechanism that makes nisin so effective as an antimicrobial agent.^6^

**Figure 1:**
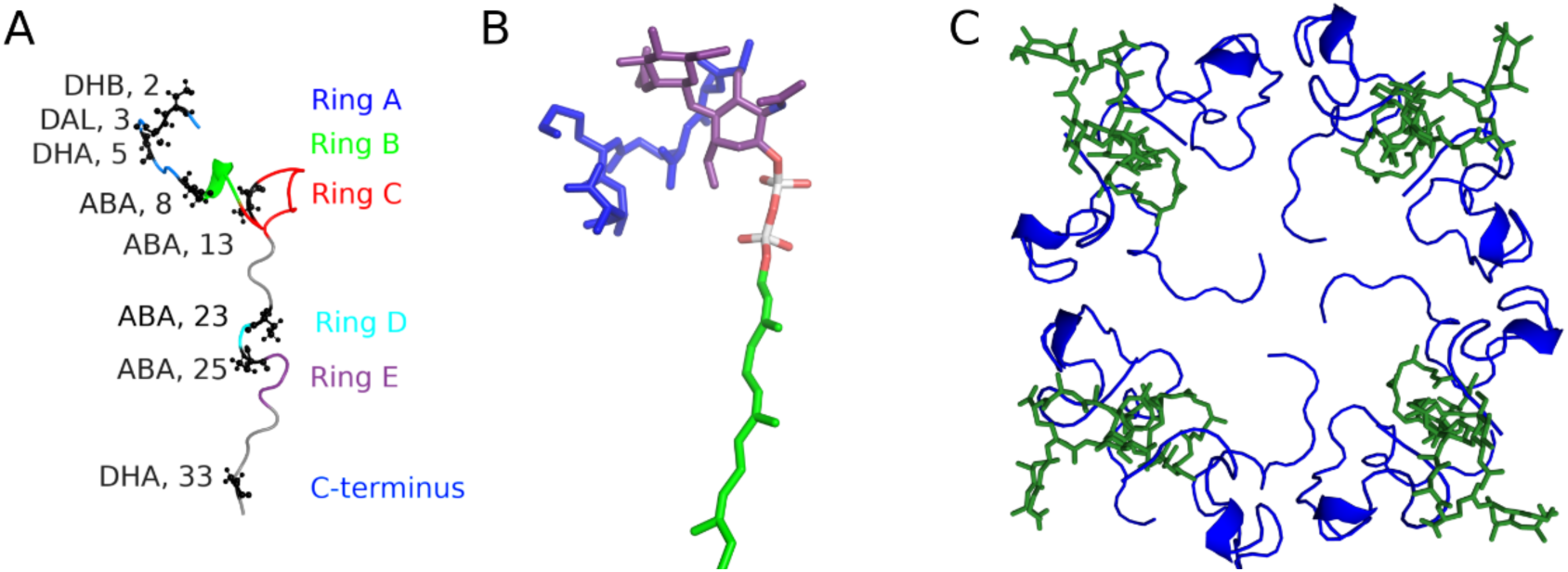
Sketch of (a) a nisin monomer, with the positions of ring A to E and the C-terminus color-coded. Listed are also the positions of the uncommon amino acids. In (b), we show the sketch of a lipid II chain, with the pyrophosphate moiety colored in red and white, the prenyl chain in green, and the additional pentapeptide and two sugars in blue and purple, respectively. In (c), we show our pore model formed by eight nisin chains (blue) interacting with four lipid II chains (green).

Despite its antimicrobial activity and being not toxic to humans, nisin is not currently used as an antibiotic. A reason for this is that at physiological pH nisin lacks solubility and stability.^7^ Attempts to design mutants that alleviate these hindrances^8–10^ have not yet led to successful candidates for new antibiotics.^11^ This could be because many of the proposed mutations focus on the flexible hinge region, and therefore may interfere with the association of nisin chains into pores perforating the cell membrane. As the process of assmbling into a pore is not understood in detail for nisin, we have carried out large-scale molecular dynamics simulations to probe the effect of lipid II in the cell membrane of gram-positive bacteria on the stability of pores formed by the nisin chains; and how the interaction between lipid II and the nisin pore changes the membrane integrity resulting in cell death.

Such computational investigations are hampered by two problems. The dehydroamino acids and thioether bridges present in nisin have not been parametrized in standard force fields. Hence, while we use CHARMM force field in our study, the force field needs to be modified to account for the uncommon amino acids. Suitable parameters were proposed in ^12,13^; and having been tested by us in earlier work,^10^ let us adjust the CHARMM36 force field as suggested by Turpin *et al.*^12^ Our modified parameters, together with our representation and parametrization of lipid II (a complex of lipids, sugars and a pentapeptide) in CHARMM36, are listed as **supplemental material**. We remark, however, that there is still an ongoing search for more effective parameters for the simulation of nisin and similar molecules.^14^

The second problem hindering our computational investigations of nisin is the large size of the system. Including thousands of water molecules, a lipid bilayer model of a cell membrane, eight nisin peptides, and four lipid II molecules, the computational complexity of the simulations is substantial. As a result, simulating the pore formation process – which in real time occurs over hours – using all-atom molecular dynamics becomes impractical. Therefore, we focused on the stability of the pore and membrane, comparing simulations of a pre-formed pore made of eight nisin chains embedded in a membrane where lipid II (characteristic for bacterial membranes) is present to that of a pre-formed pore in the absence of lipid II. Our analysis, covering more than 200 ns per trajectory, indicates that lipid II increases not only the stability of the nisin pore, but that its interaction with nisin also leads to local changes in thickness and viscosity within bacterial membranes, ultimately weakening their integrity.

## II. MATERIALS AND METHODS

### A. System Preparation

Our simulations, comparing the stability of a pore built from nisin-chains in the presence and absence of lipid II, rely on three elements: generation of a self-actualized membrane, the design and positioning of the nisin-pore complex into the membrane, and the addition and positioning of lipid II molecules.

#### Membrane Bilayer

As nisin is effective against gram-positive bacteria and therefore interacts with gram-positive membranes, we used the CHARMM-GUI Membrane builder^15^ to generate a realistic model of a gram-positive bacteria cell wall with a 3:1 ratio of 1-palmitoyl-2-oleoyl-sn-glycero-3-phosphoglycerol (POPG) with 1-palmitoyl-2-oleoyl-sn-glycero-3- phosphoethanolamine (POPE))^16,17^, *i.e.*, 528 POPG and 177 POPE lipid molecules. Our membrane bilayer model was placed in a 150 x 150 x 230 Å box and was solvated with 137,413 water molecules. Following the CHARMM-GUI membrane builder ion option, we neutralize the system with 0.15 mM KCl. The resulting membrane model was first minimized for 5000 steps and afterward simulated in short molecular dynamic simulations at 300 K and constant volume, gradually increasing step size and step time according to the standard CHARMM protocol^15,18–20^. To equilibrize the lipid motions and minimize flip-flops of the bilayer lipid molecules, we simulated the membrane for an additional ∼1 microsecond at a constant temperature and pressure of 300 K and 1 bar. Finally, we removed the water and ions in this self-actualized membrane, leaving only the bilayer to build the systems considered for our simulations.

#### Nisin-Pore Complex

In the second step, we inserted a pore complex made of eight nisin peptides and four lipid II molecules in the so-generated self-actualized membrane. The pore was designed to have a diameter of 25 Å, enclosed by eight nisin molecules arranged in a circular fashion, with each lipid II molecule situated around two nisin molecules. The number of nisin proteins forming the pore and the pore size reflect the available experimental data. Breukink et al. used pyrene fluorescence and CD to show that the lipid II and nisin chains are arranged in the membrane bilayer in an 8:4 ratio.^21^ Sahl *et al.* showed, through electrochemistry, that this ratio is constant and that the diameter is ∼20-25 Å.^22^ As Medeiros *et al.* pointed out, residue 28 is the last residue in the membrane,^23^ and the disordered C-terminus is outside of the membrane. Taking this into account, the nisin chains in our pore model must be long enough to span both layers of the membrane. However, the sole model of nisin bound to lipid II, resolved by solution NMR and deposited in the Protein Data Bank (PDB) under identifier 1WCO, measures only 30 Å between the N-terminus and residue 28, while the bilayer has a thickness of about ∼43 Å. To obtain sufficient long nisin chains, we have simulated the nisin chains (starting from the 1WCO conformation) for one ns at 315 K, allowing them to unwind partially. The resulting conformation is still similar to the 1WCO model, but with a length of 40 Å between the N-terminus and residue 28 fits now to the thickness of the bilayer. This structure also has residue 17 sitting in the middle of the bilayer as indicated by both Medeiros^23^ and Breukink.^24^

Experimental data were also used to position the lipid II molecules. For instance, Ganchev et al. showed (using AFM imaging) that the pentapeptide located at the head group of lipid II is just exterior to the bilayer surface and points away from it,^25^ and Chugnov et al.^16^ showed that the pyrophosphate moiety of lipid II are similarly placed as the phosphate groups in the POPE and POPG headgroups in the bilayer. For this reason, we placed the lipid II head group in such a way that their phosphorus atoms are adjacent to the phosphorus groups of other lipids in the bilayer. As the Dodecaprenyl-C55 chain is highly conserved in bacteria, a chain of 55 carbons was used^23^ as the acyl chains of lipid II. While these lipid II chains can adopt either an L, I, or V shape^16^, only the I-shape allows them to form contacts with the nisin peptides. Therefore, we considered only this shape, a choice that is justified, as Chugnov et al. also showed that interactions between the head group and nisin are more important.^16^ As the N-terminal residues are known to bind to the lipid II molecules through hydrogen bonding involving the amino hydrogens in the backbone of the PPI group^23^, we added the lipid II molecules at a distance of around ∼3.5 Å to the first four residues of nisin.

#### Insertion of the Pore into the Membrane

The above-generated pore complex of nisin and lipid II molecules was then placed into the self-actualized bilayer in an opening of about ∼40 x 40 Å (which exceeded the pore by about two Å) that was created using the emulate function in Pymol. The resulting membrane-pore system was solvated in 40,000 TIP3P water molecules and 0.15 mM NaCl in a box 150 x 150 x 80 Å. The resulting model was again minimized over 5000 steps and simulated over 1.5 ns at a constant volume and temperature of 315 K. For equilibration, we carried our molecular dynamics runs at 315 K and one bar but restrained the pore in an initial step for five ns with a factor of 1000 kJ/mol. In three additional steps of five ns each, these constraints were gradually removed in steps of 250 kJ/mol. This procedure leads to an equilibrated conformation that was used as the starting point for our simulations of nisin pore interacting with lipid II and the lipid bilayer in our model of a bacterial cell membrane.

Using Pymol’s atom selection feature to remove the lipid II molecules in the above model and simulating the resulting system with restrained positions of the nisin atoms for another one ns, we later generated a control model for the membrane-pore system. Lacking lipid II molecules, this control allows us to pinpoint how lipid II affects the stability of the nisin pore and its interaction with the bilayer.

### B. General Simulation Protocol

The stability of the above-designed nisin-pore-membrane systems in absence and presence of the bacterial membrane-specific lipid II are studied by molecular dynamic simulations utilizing the GROMACS 2022 and 2023 software packages.^26^

As nisin contains uncommon amino acids (dehydroalanine and dehydrobutyrine), and one lanthionine and four β-methyllanthionine), we had to modify the CHARMM36 all-atom forcefield^27^ used by us in combination with TIP3P water^28^ to describe inter and intramolecular interactions. Similarly, we had to derive topology and parameters for the lipid II molecules that are shown in the **Figure 1b**. Eleven isoprene units, the first seven units in the (Z)-configuration followed by three in the (E)-configuration, form a lipophilic anchor.^29–32^ It is connected by a pyrophosphate group to the polar headgroup made of a N-acetylglucosamine sugar (Glc) which is attached to a N-acetylmuramic acid sugar (Mur). Additionally linked to Glc is a peptide whose composition varies depending on bacterial species; commonly employed in research studies and also used by us in this study is the L-Ala–γ-D-Glu–L-Lys–D-Ala–D-Ala pentapeptide.^16,33^ The saccharide MurNAc was created by combining the topologies of N-acetylglucosamine and 2-methoxy-propionate, both from the CHARMM36 carbohydrate force field.^34^ Parameters for the non-proteinogenic peptide linkages in lipid II were taken from the CHARMM36 protein force field^27^ and partial charges were assigned to resemble that of the peptide linkage in proteins. The additional parameter and topology files can be found in the **supplemental material**.

The membrane-pore complex was solvated by adding water such that the box is extended in z-direction on both sides by 20 Å. We then added 0.15 M NaCl ions to neutralize the system. Periodic boundary conditions are employed in all directions, and long-range electrostatic interactions are calculated with the particle-mesh Ewald (PME) technique using a real-space cutoff of 12 Å. Short-range van der Waal interactions are truncated at 12 Å with smoothing starting at 10.5 Å. Before the production run, each system was minimized with 50,000 steps with the steepest descent integrator in^26^ and then equilibrated at constant volume for 750 ps at a temperature of 315 K with a leap-frog integrator^26^ and a step size of 0.001 fs and then again for an additional 750 ps with a step size of 0.002 fs.

After equilibration, each system (the one which has lipid II interacting with the pore and membrane is named by us e*xperiment*, and the other where lipid II is absent we termed *control*) was followed in three independent trajectories that differ their initial distribution of velocities. The lengths of the six trajectories are listed in **Table 1**, with in each simulation the temperature and pressure set to 315 K and one bar, as controlled by a Nose-Hover thermostat^35,36^ and Parrinello-Rahman barostat.^37^ Using the SETTLE^38^ and LINCS^39^ algorithms to control fast fluctuations and to restrain hydrogens allows us to use a time step of 2 fs for integrating the equations of motion. PDB files of start and final conformations are available as **supplemental material**.

**TABLE 1:** System Set-up.

| System | # Atoms | #water molecules | #Independent Trajectories | Trajectory length (ns) | Total sampling (ns) |
| --- | --- | --- | --- | --- | --- |
| Control<br>(w/o lipid II) | 183,347 | 30,968 | 3 | 200 | 600 |
| Experiment<br>(with lipid II ) | 184,479 | 30,968 | 3 | 200 | 600 |

### C. Trajectory Analysis

The molecular dynamics trajectories are analyzed with the GROMACS tools^26^, VMD^40^, MDAnalysis^41,42^, and Membrainy^43^. For visualization of trajectory conformations, we use VMD. Common quantities such as the root mean square deviation (RMSD) or root mean square fluctuation (RMSF) are calculated using GROMACS tools, using for the solvent-accessible surface area (SASA) a spherical probe of a 1.4 Å radius. Contact frequencies are computed with VMD, where we define contacts by a cutoff of 4.5 Å in the closest distance between heavy atoms in a residue pair. Hydrogen bonds are defined by a distance cutoff of 3.5 Å between the donor and acceptor atoms and an angle cutoff of 30°. Membrane properties such as membrane thickness area per lipid, head group angles, and chain order parameters were calculated with Membrainy, with the respective molecules selected by MDAnalysis when calculated only for a shell at a certain distance to the pore.

## II. RESULTS AND DISCUSSION

### A. Lipid II modulated Nisin Pore stability

We start with looking into the stability of the pore, built from eight nisin peptides and an additional four lipid II molecules, inserted into a bilayer formed from POPE and POPG lipids. Data from three independent trajectories (the *experiments*) covering 200 ns are compared with three trajectories of the *control* (where lipid II is absent) having the same length. In **Figure 2**, we contrast for each trajectory the final conformation with the start conformation. For both the experiment and the control systems, we see a slow decay of the pore, but the change is smaller in the three trajectories where lipid II interacts with the nisin pore.

**Figure 2:**
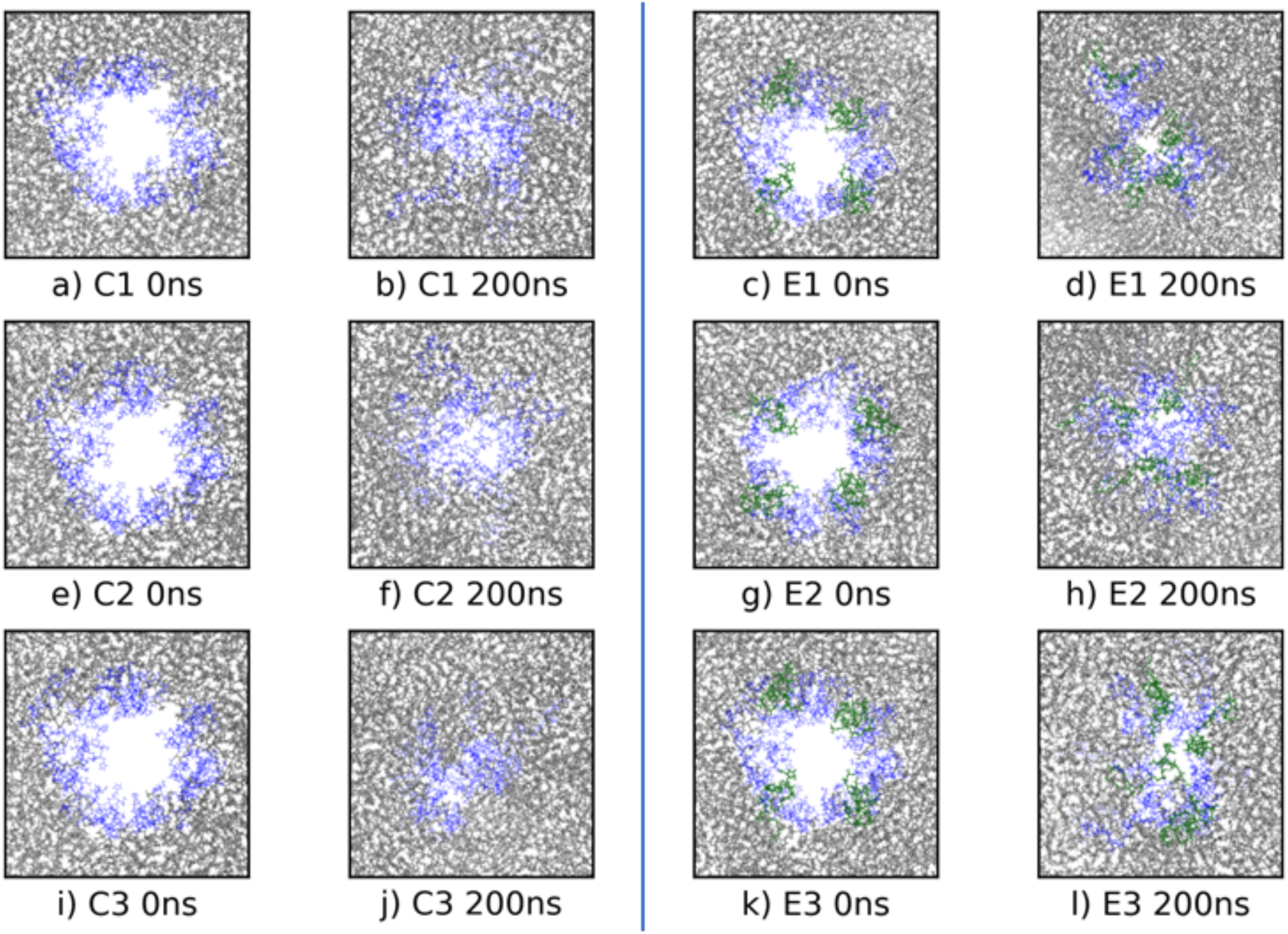
Start and final conformations of six trajectories of the nisin pore embedded in the membrane, either in absence (the control, C1 - C3), and presence of lipid II (the experiment, E1-E3). Nisin chains are colored in blue while lipid II molecules are drawn in green, and POPE or POPG lipids in gray.

To quantify the decay of the pore, one needs to define and measure its extension. Because of the irregular shape of the pore, we chose for this purpose as a proxy the amount of water inside the pore. To account for the dynamically changing thickness of the membrane, we first calculated the center of mass of all eight nisin chains, taking into account only residues 8-28 located within the bilayer. We then measured the distance to the center of mass for each of the eight chains and chose the largest distance as the radius for a sphere around the center of mass. In this sphere, we deleted all water molecules that are outside the bilayer and the nisin assembly, counting only the remaining water molecules. The numbers obtained, as measured in the start and final conformations, are listed in **Table 2**. Note that in run E1, not only did one of the nisin chains detach from the pore, but also one of the lipid II molecules lost contact with the pore and drifted away, which explains why this trajectory led to a more significant loss of water molecules and is indicative of the significance of the stereochemistry and presence of the lipid II molecules.

**Table 2:**
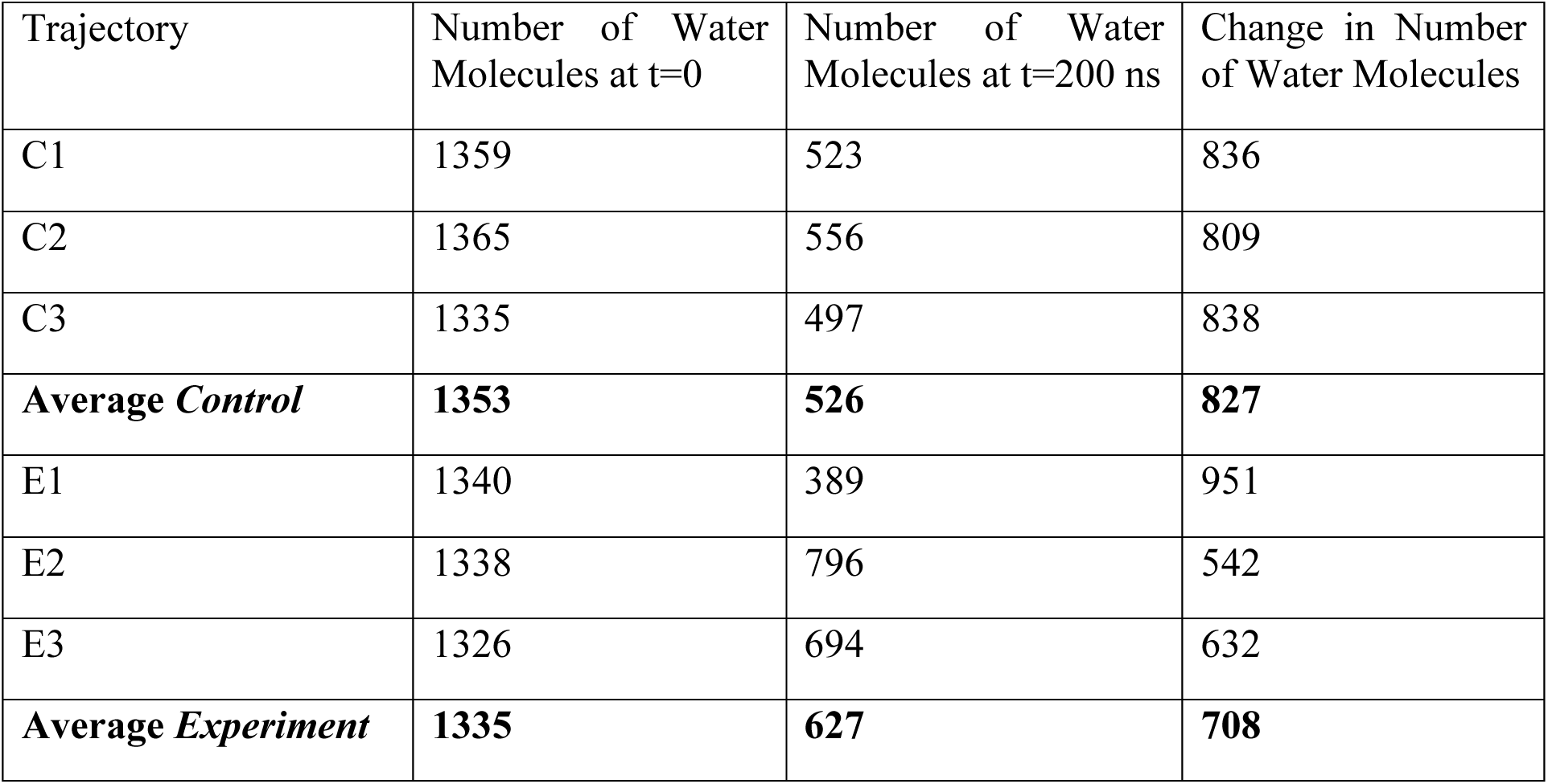
Number of water molecules inside of the nisin pore assembly for both control (lipid II missing) and experiment (where lipid II interacts with the nisin chains). Data are shown for the start and the final frame, and their difference.

Comparing the initial and final conformations, we see that the number of water molecules at the start is similar in all six runs. However, at the end of the trajectories, the number of water molecules differs substantially between the control and experiment simulations. On average, one finds in the experiment trajectories approximately 100 additional water molecules within the pore, indicating a difference in the pore volume of about 1150 Å^3^; at the end of the trajectory, the control has an average pore volume of 6000 Å^3^, and the experiment has an average pore volume of 7150 Å^3^. Assuming a membrane thickness of about 30 Å and cylinder geometry for the pore, this change in pore volume corresponds to a pore diameter that is around 1 Å larger in diameter, i.e., the diameter of the pore in the presence of lipid II is 9 Å compared to 8 Å in the control where lipid II is missing.

In order to describe the evolution of the pore with respect to time in more detail, we have analyzed the average root-mean-square-deviation (RMSD) with respect to the start conformation as a function of time (see **Figure 3**). In **Figures 3a and 3b**, the RMSD is evaluated over the whole pore-forming assembly of nisin chains. We notice that this global RMSD does not depend on whether we consider for the RMSD calculation only the backbone Cα-atoms (**3a**) or all heavy atoms (**3b**) of the nisin chains, with differences in the plots barely noticeable. This indicates that the overall change in the pore geometry is driven by the re-arrangement of the complete nisin chains and not dominated by side-chain motion. When lipid II is present, the RMSD values are slightly lower, especially at the early stages of the simulation, and have a smaller standard deviation. However, after about 125 ns, the RMSD plots for both the control and the experiment plateau at similar values and with comparable standard deviations. The similarity in RMSD is reflected by similar values in the number of inter-chain contacts measured between the nisin molecules: 93(16) in the experiment (the bacterial membrane model containing lipid II) vs 118 (11) in the control. Note that none of these contacts is present in the start conformation; hence, while we find in both cases an increase of about 60 interchain contacts (72 in the presence of lipid II and 56 in the control), the larger number of contacts results from a decay of the pore that allows for movement of the nisin chains.

**Figure 3:**
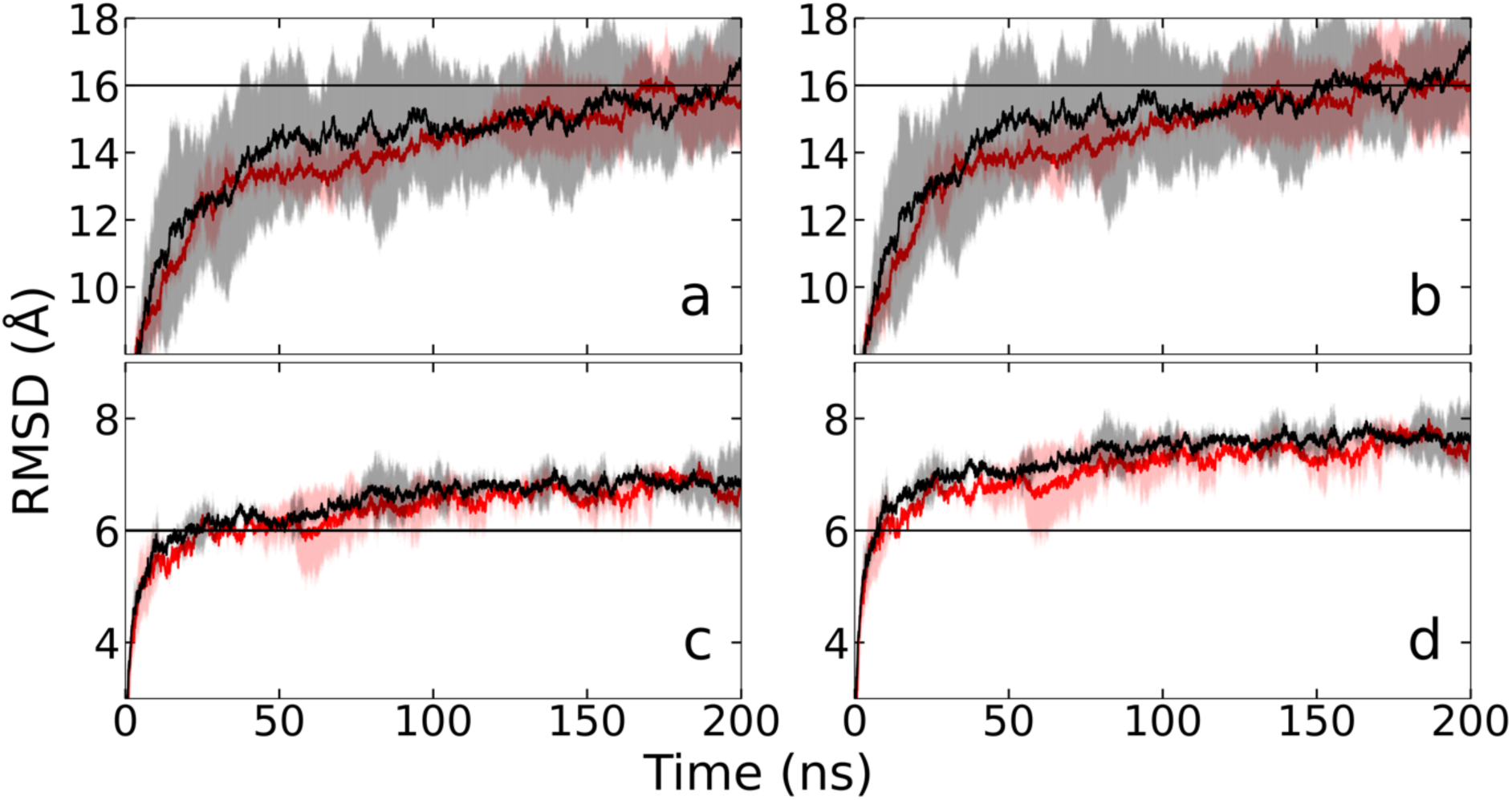
Global Root-mean-square deviation (RMSD) with respect to the start conformation as a function of time for the nisin assembly in the presence (Experiment, shown in red) and absence (Control, shown in black) of lipid II, calculated over Cα-atoms (a) and all heavy atoms (b). The corresponding plots for the chain RMSD are shown in (c) and (d). Horizontal lines are drawn to guide the eyes and to emphasize differences between the sub-figures.

A different picture is seen in **Figures 3c** and **3d**, where the RMSD is now evaluated separately for each of the eight chains, with the average over the eight chains (and the three trajectories) shown as a function of time. In these two sub-figures, we see that the chain RMSD values are larger when calculated over all heavy atoms (**3d**) instead of only Cα-atoms (**3c**), indicating that changes in the side chain orientations contribute strongly to the chain RMSD. Note that the standard deviations for both the lipid II system and the control are smaller than for the global RMSD plots in **Figures 3a** and **3b**. Unlike the global RMSD, chain RMSD values for the experiment are lower than the control over most of the trajectory, with the difference appearing to be also more pronounced. While the number of intra-chain contacts (connecting residues within the same chain) stays for both control (about 23 contacts) and in the experiment (about 22 contacts) unchanged throughout the simulation, this is not true for individual chains, where contacts are both lost and newly formed. Especially the large standard deviations in the chain RMSD for the lipid II system seen between 50 ns and 125 ns (and again 150 ns -175 ns) indicate significant variations in the chain geometries between the independent trajectories. However, all experiment trajectories still have smaller chain RMSD than seen in the trajectories of the control.

The visual inspection of the final conformations of our six trajectories and the estimated change in pore volume indicated that lipid II in the membrane stabilizes the pore-like nisin assembly. The four RMSD plots in **Figure 3** not only support this observation but also suggest that lipid II stabilizes the nisin pore mainly by preserving the fold of the nisin peptides in the pore, thereby slowing down the decay of the pore. In order to establish the regions in the nisin chain conformations that are affected most by the presence of lipid II, we show in **Figure 4** the root-means-square fluctuation (RMSF) of residues averaged over all eight chains, a quantity that allows one to identify the regions in the nisin chains that are most flexible and prone to change. The RMSF is evaluated with respect to the start conformations, but we remark that the overall picture does not change when other reference conformations (for instance, such as taken from the mid-points of the trajectories) are chosen (data not shown).

**Figure 4:**
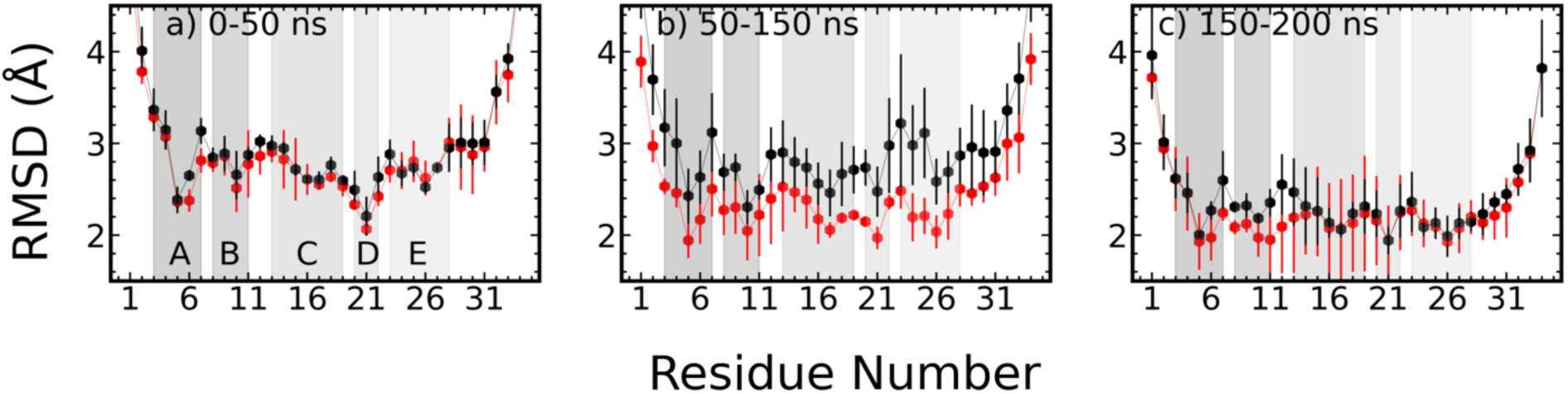
Root-mean-square fluctuation (RMSF) of nisin residues (averaged over all eight nisin chains in all three trajectories) in the presence (Experiment, drawn in red) and absence (Control, drawn in black) of lipid II. In (a), we show the values calculated over the first 50 ns; in (b), the values measured between 150-150 ns, and in (c), the ones obtained for the final 50 ns. Shaded areas mark the residue regions corresponding to the five rings A to E.

When evaluated over the first 50 ns (**Figure 4a**), the RMSF values in the absence and presence of lipid II differ little and agree within the error bars. This is different for the interval 50ns-150ns (**Figure 4b**), roughly corresponding to the time span, whereas in **Figures 3c and 3d**, the chain RMSD has a large standard deviation in the presence of lipid II. Here, we observe a pronounced lowering in the residue-wise fluctuations over the regions of residues 8-28 (which are embedded in the membrane) in the experiment (i.e., when lipid II is present), while there is little change in the values for the control. These differences start at the hinge region and ring D (residues 21-28) and around residues 8-11, i.e., ring B of the nisin chain (see **Figure 1a**), before later extending over all membrane-embedded residues. Over the last 50 ns (see **Figure 4c**), the differences between the control and the pore in the presence of lipid II decrease as the fluctuations in the control decrease, with the remaining differences again most pronounced for residues 8-13, i.e., ring B and residues 12 and 13 that lie between ring B and C and are important for the flexibility of the chain. Mutation in this residue has been shown to have enhanced antimicrobial activity^44^.

In order to understand what causes the observed differences in the mobility of the nisin residues when interacting with lipid II, we list in **Table 3**, the solvent-accessible surface area (SASA) measured for these segments both at the start of the trajectory and averaged over the last 50 ns of a trajectory. The listed SASA values describe the area of nisin chains in the pore that is exposed to the outside (i.e., the expanse of the pore) and not covered by intra-chain contacts, i.e., contacts with other nisin-chains or with lipid II molecules. Hence, it describes the area available for contact with either water molecules or POPE and POPG lipids in the membrane. By excluding the area covered through contact with POPE and POPG lipids, we arrive at the water-exposed area WASA, also listed in **Table 3**. The difference between the two quantities is the surface area covered by POPE or POPG, named by us LASA and again listed in **Table 3**. Note that we also list for all three quantities the difference between initial values and the averages over the final 50 ns, which quantifies for the segment change over the trajectory.

**Table 3:** Change in solvent accessible surface area (SASA) between the last 50 ns and the respective start conformation. SASA can be exposed to either water or lipids (POPE and POPG). If the area covered by POPE or POPG is excluded, we call the resulting quantity water-accessible surface area (WASA). In contrast, when they are exposed to POPE and POPG, we call it lipid-accessible surface area (LASA). Values are given in Å^2^ and reported for both the experiment (presence of lipid II) and control (absence of lipid II). We also list the differences between the two systems (ΔΔSASA and ΔΔWASA). Our data are evaluated for different segments of the nisin chains, with residues 8-28 inside the bilayer and residues 1-7 and 29-34 outside the membrane. The standard deviation of the averages is given in parentheses, marking changes for the last digit.

| Residues | $\Delta\text{SASA}$ | $\Delta\text{WASA}$ | $\Delta\text{LASA}$ | $\Delta\Delta\text{SASA}$ | $\Delta\Delta\text{WASA}$ |
| --- | --- | --- | --- | --- | --- |
|  | E \| C | E \| C | E \| C |  |  |
| 1-34 | -1.3(1.1) \| -2.6(4) | -4.2(4) \| -4.2(5) | 2.9(1.2) \| 1.5(9) | 1.3(1.1) | -0.01(50) |
| 1-7 | 0.6(6) \| -0.8(2) | -1.6(1) \| -1.8(1) | 2.1(8) \| 1.0(5) | 1.4(5) | 0.2(2) |
| 8-11 | -0.1(2) \| -0.3(1) | -0.2(3) \| -0.2(2) | 0.1(6) \| -0.1(5) | 0.1(2) | 0.02(40) |
| 8-28 | -1.6(7) \| -1.5(4) | -1.8(3) \| -2.0(4) | 0.2(1) \| 0.5(9) | -0.1(5) | 0.2(1) |
| 12-20 | -1.1(4) \| -0.78(3) | -1.0(1) \| -0.8(1) | -0.1(7) \| 0.04(60) | -0.3(4) | -0.2(2) |
| 21-28 | -0.3(3) \| -0.5(2) | -0.6(4) \| -0.9(4) | 0.3(8) \| 0.4(8) | 0.2(1) | 0.3(4) |
| 29-34 | -0.3(3) \| -0.3(1) | -0.8(2) \| -0.3(3) | 0.5(7) \| 0.04(60) | 0.02(20) | -0.5(4) |

We see that independent of the absence or presence of lipid II, the pore-forming nisin molecules lose accessible surface area over the course of the simulations; however, the loss of WASA is more significant than that of SASA, i.e., the surface area that at the start was exposed to water is now exposed to POPE or POPG, pointing to the bilayer. Hence, the difference between SASA and WASA describes the loss of pore geometry and the succeeding mixing of nisin molecules with the bilayer. Over the whole length of the nisin chain, the area covered by POPE and POPG in the presence of lipid II increases by 2.9(1.2) nm^2^, while in the control by only 1.5(9) nm^2^. The difference arises mainly from residues 1-7, which interact with the lipid II molecules: an additional area of 2.1(8) nm^2^ versus 1.0(5) nm^2^ in the control. For residues 8-28, which are embedded in the bilayer, the area that is exposed to the bilayer (i.e., POPE and POPG molecules) increases by 0.5(9) nm^2^ in the control but only by 0.2(1) nm^2^ for the experiment, i.e., in presence of lipid II. This indicates that in presence of lipid II the side chains of nisin are more oriented into the pore and therefore interact less with the bilayer, stabilizing in this way the pore.

What causes the increased stability of the nisin pore in the presence of lipid II molecules? One factor may be the contacts between the nisin chains and the lipid II molecules and their reorganization along the trajectory. At the start, the eight nisin chains share a total of 45 contacts with the four lipid II molecules, with each lipid II forming contacts with two neighboring nisin chains, around 5-6 contacts with each chain. These contacts are primarily with N-terminal residues of the nisin chains, about three with the mostly extracellular residues 1-7 (mostly with the first three residues). About two contacts are with residues 8-28, which are located inside of the membrane bilayer. At the end of the trajectory, the four lipid II molecules still form about 38 contacts with the nisin chains; however, each lipid II has now, on average, only about two contacts with the exterior segment (residues 1-7) of a nisin chain, but has increased the number of contacts with the interior residues 8-28 of a nisin chain by around 2-3 contacts. These contacts, formed mainly by the head group atoms of lipid II with adjacent nisin molecules, provide additional stabilization to the pore geometry. Note that these contacts involve the pyrophosphate moiety but not the pentapeptide of lipid II, i.e., binding of nisin to lipid II cannot be evaded by mutations in the pentapeptide. This is similar to what is observed for another antimicrobial peptide, teixobactin^45^, and may explain why resistance to nisin is rare. Note also that these contacts coordinate the lipid II head groups to nisin chains. Visual inspection of frames along the trajectories E1, E2, and E3 indicates that the resulting proximity of the flexible and moving lipid II tails constrains the movement of the nisin molecules within the membrane, therefore adding to the stability of the pore. However, as discussed by us later in the following section, this movement of the lipid II tails (with a RMSF of about 6 Å for the final ten carbon atoms) also disturbs and weakens the membrane directly. Interestingly, we did not see that lipid II changes the interaction of Na^+^ ions with nisin and the lipid bilayer as was observed in previous work^46^. Specifically, we did not see any specific change in the binding pattern of the ions between control and experiment, and therefore, do not believe that such ion-mediated interactions contribute to the stability of the pore.

### B. Nisin - lipid II Mediated Membrane Integrity

What are the effects of the nisin pore complex on membrane integrity? The change in the surface area of the nisin chains that are exposed to either water or POPE/POPG lipids leads to a change in the number of contacts and hydrogen bonds between water or POPE/POPG molecules with nisin chains. We show this for water molecules in **Figures 5a** and **5c**. For residues 8-28 (i.e., the ones embedded in the interior of the membrane), the number of hydrogen bonds is reduced by 40 in the control (i.e., about five hydrogen bonds per nisin chain) but only by around 30 in the experiment, in presence of lipid II (about four hydrogen bonds per chain). Approximately 16 hydrogen bonds (two per chain) are lost for both systems in the region of residues 12-20. On the other hand, the total number of contacts between water and nisin chains is for residues 8-28 about 40 contacts lower in the presence of lipid II than in the control, with the difference mainly resulting from residues 12-20. This is because in the control, some nisin residues no longer form hydrogen bonds with water molecules, but they stay close enough to the water molecules to form contacts, i.e., the lost hydrogen bonds are now counted as contacts. Nevertheless, contact and hydrogen bond numbers together indicate that the presence of lipid II stabilizes the hydrogen bonds between water molecules and the nisin residues 8-11 and 21-28, which corresponds to ring B and ring D. As a result, the two rings are more likely oriented towards the water, pointing towards the pore interior thus constraining and stabilizing the pore geometry.

**Figure 5:**
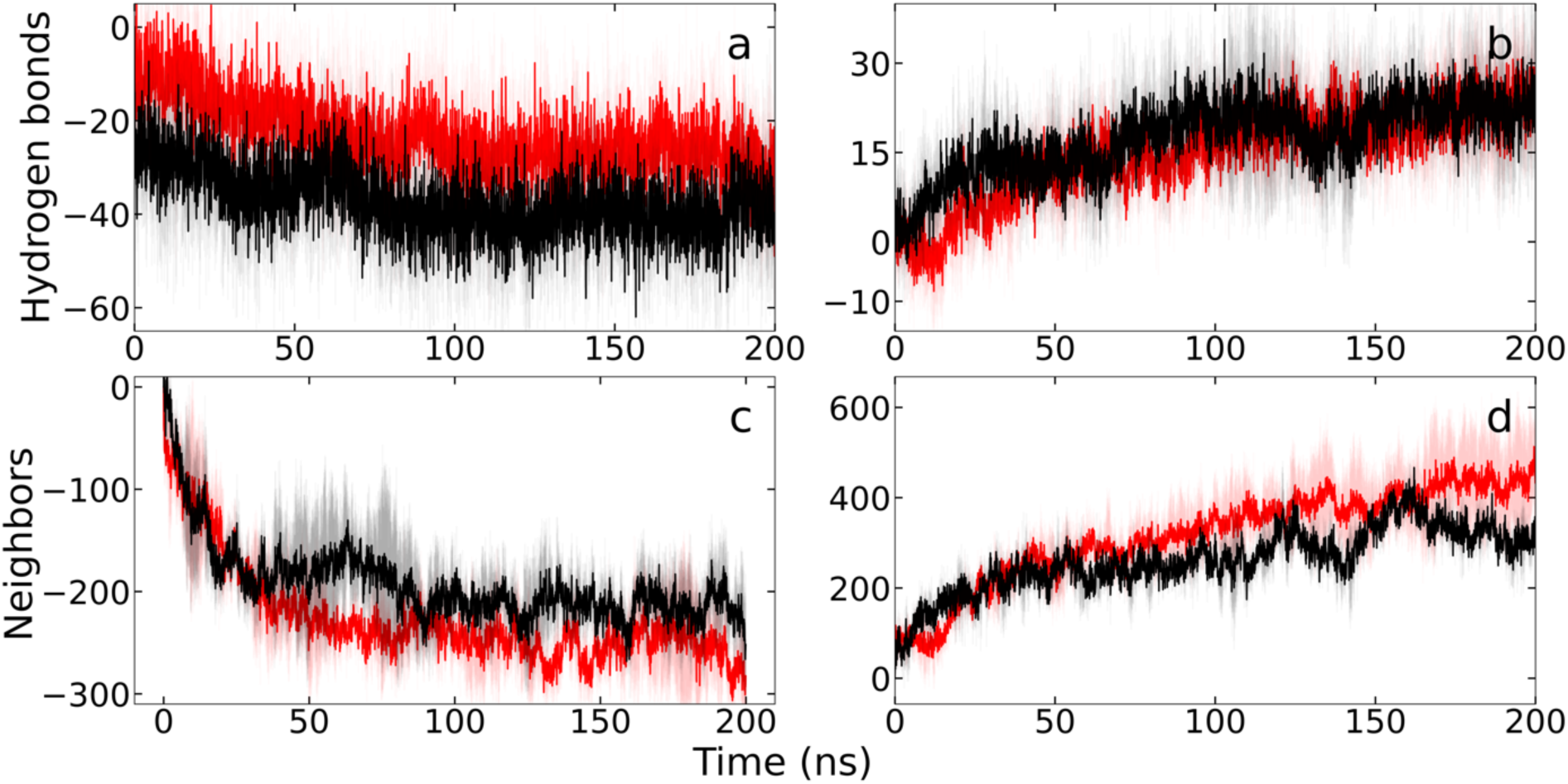
The number of hydrogen bonds formed between nisin chains and water molecules located within the pore are shown in (a). Only the residues 8-28, located in the interior of the membrane, are considered. The plots show the difference between the average, taken over the last 50 ns and three trajectories, and the initial values, i.e., a negative value indicates a loss of hydrogen bonds. Similarly, we show in (b) the corresponding number of hydrogen bonds formed with either POPE or POPG lipids. The number of water molecules that do not form hydrogen bonds with nisin residues but are within 4.5 Å to nisin residues is shown in (c), and the corresponding number for contacts between nisin chains and POPE or POPG is shown in (d). Red indicates the experiment, and black the control.

Residues in the nisin chains can also form contacts with the POPE and POPG lipids that make up the membrane bilayer. The time evolution of these contacts is shown in **Figures 5c** and **5d**. The number of contacts between residues 8-28, embedded inside of the membrane, and POPE lipids on average decreased by around 20 contacts for the control group: at the beginning of the trajectory, there were 121 contacts which fell to an average of 104(14) contacts in the last 50 ns of the trajectory. However, in presence of lipid II, the number of contacts between residues 8-28 and POPE lipids on average increased by around 40 contacts; at the beginning of the trajectory there were 93 contacts which increased to an average of 125(4) contacts in the last 50 ns, with almost 30 contacts formed by residues 8-11 (ring B) and 21-28 (ring D), see **Figure 1**. The corresponding number of contacts between the nisin chains and the more frequent POPG molecules increases for the experiment, i.e., in the presence of lipid II, by about 170 contacts, beginning with 394 contacts at the start of the trajectory and increasing to an average of 568(72) contacts in the last 50 ns. However, the increase is larger in the control, where the number of contacts grows by about 190 contacts, initially having 412 contacts, which increased to an average of 600(60) contacts at the end of the trajectory. Again, most of the newly formed contacts of POPG are with nisin residues 8-11 (ring B) and 21-28 (ring D): about 90 in the experiment and 100 in the control.

Hydrogen bonds between the nisin chains and POPE are rare and do not change in frequency; on average, we observe, during the last 50 ns, 3(1) hydrogen bonds in the control and 4(2) hydrogen bonds in the presence of lipid II, i.e., in the experiment. On the other hand, hydrogen bonds are not only seen, but increase, between the nisin chains and the more numerous POPG: there was an increase of 13 hydrogen bonds for the control (beginning with 13 hydrogen bonds and increasing to 26(5) at end of the trajectory) and an increase of around 9 hydrogen bonds for the experiment (beginning with 17 hydrogen bonds and increasing to 26(4) at the end of the trajectory). Note that about 66% of the new hydrogen bonds are formed between POPG and residues 8-11 or 21-28: 9 in the control and 6 in the experiment.

Hence, both the number of hydrogen bonds and the total number of contacts between the nisin chains and the POPE/POPG lipids increase over the lengths of the trajectories, but less so in presence of lipid II. This is consistent with the stronger decay of the pore in the control, leading to more nisin residues coming in contact with the membrane lipids while reducing the number of hydrogen bonds formed with water. Most of this effect comes from rings B (residues 8-11) and D (residues 21-28). The contact of these residues in the nisin chains with lipid II encourages an orientation of these rings where they face toward the pore, which stabilizes hydrogen bonds with water molecules and reduces the chance for contacts or hydrogen bonds to form with the membrane constituents, i.e., the POPG/POPE lipids. This explains the differences between our experiments and the control runs.

This growing interaction of the nisin chains with POPE and POPG molecules, as seen throughout the respective trajectories, disturbs the membrane. At the start of the simulations, there are no significant differences in the distribution of lipids around the pore between the control and the experiment. This is different at the end of the simulations, when, in the vicinity of the pore (defined as the shell with a radius between 25 Å and 40 Å from the center of the pore), the number of POPE and POPG molecules increases from 113 to 141 molecules in the control, but only from 109 to 134 molecules in the experiment, i.e., in presence of lipid II. Hence, while at the start, the density of lipids was similar in the control and experiment, it became lower in the experiment than in the control as the simulation evolved. This is because the pore decays in both experiment and control over the course of the trajectories, allowing for an inflow of POPG and POPE lipids. This influx is more pronounced in the control than in the experiment where presence of lipid II limits this effect. Correspondingly, there is a difference in the thickness of the membrane in the vicinity of the nisin assembly: a membrane thickness of 31.8(8) Å is measured in the control but only 27.5(1) Å in the experiment. For distances larger than 40 Å, the densities and thickness of the bilayer (on average 30.4(1) Å over the last 50 ns in both systems) are again similar in experiment and control and changed little between the start and finish of the trajectories. This indicates that the presence of lipid II not only stabilizes the nisin pore but also leads locally to a thinning of the bilayer.

One way to characterize the behavior of the membrane is by the area per lipid (APL), This quantity is defined as the area of a selected segment (“box”) of the membrane divided by the number of lipid molecules in the box. As this quantity changes in response to the presence of proteins and other non-lipids, we have measured it for both the control and the experiment trajectories as a function of distance to the center of the pore. The APL was evaluated separately for the two layers of the membrane as the lipid II molecules are located in the layer close to the extracellular space and the N-terminus of the nisin chains. In our notation, this is layer 1. In bulk, the difference between the two layers is neglectable in both control and experiment; however, differences are seen near the pore. Comparing the averages taken over the last 50ns and all three trajectories of each system, we notice that in the shell with a radius between 25 Å and 40 Å from the center of the pore, the APL is in both layers larger in the presence of lipid II than in control, and in both cases are the APL values larger in layer 1 than in layer 2: in layer 1, we have in the control a value of 54.1(7) Å^2^ vs 56.0(6) Å^2^ in the experiment, while in layer 2 the corresponding numbers are 47.2(6) Å^2^ for the control and 50.5(6) Å^2^ in the experiment. The APL values for layer 1, in both the control and the experiment, remain essentially unchanged from the beginning to the end of the simulation, 53.7 Å^2^ and 55.6 Å^2^, respectively. However, the APL values decreased for layer 2 in both absence and presence of lipid II: the start value for the control is 54.7Å^2^ and 56.6 Å^2^ in the experiment. These differences of the time evolution of APL values show that the presence of the nisin chains distorts layer 2 more than layer 1 but that this effect is lesser in experiment, i.e., is restrained by presence of lipid II; in the control is in layer 2 the APL by 6.9(7) Å^2^ smaller than in layer 1, but the difference is in the experiment only -5.5(6) Å^2^. Note that at the start, the corresponding differences are about 1 Å^2^ in both the control and in the presence of lipid II.

Another way to characterize the membrane behavior is through the head group angles of the lipids in the membrane. For the head group atoms, this orientation can be described by head group angles defined such that for zero angle, the lipid head is parallel to the membrane surface, and for positive angles points away from the membrane. For the less frequent POPE in the vicinity of the pore (in a shell with a radius between 25 Å and 40 Å), we find that there are no significant differences in the distribution of head group angles when comparing the control to the experiment. Regarding the POPG headgroup angles, there is only a slight shift toward more negative headgroup angles. In all cases, the averages are compatible with zero, which suggests that the changes in membrane properties more likely result from the orientations in the lipid tails.

A measure for this quantity is the order parameter, -S_CD_, which characterizes the distribution of angles for the carbon atoms in the POPE and POPG lipids, i.e., it describes the fluidity of the lipids in a membrane segment. By its definition, this quantity takes values between -1 (disorder) and 0.5 (completely ordered)^43^. Values of this quantity, as measured and averaged over the final 50 ns, differ in both layers little for POPG and POPE lipids in bulk or far away from the nisin assembly. However, when measured in a shell of radius between 25-40 Å, i.e., close to the pore, the order parameter values are for layer 2 in the presence of lipid II higher than in the control. However, for layer 1, where the peptide chains of lipid II reside, the values are lower than in the control. This can be seen in **Figure 6**, where we show either for each layer the differences between order parameters measured in experiment trajectories and the ones measured in control runs (in Figure **6c** for POPE and in **6d** for POPG), or the differences between order parameter values measured in the two layers (in Figure **6a** for POPE and in **6b** for POPG) in either experiment or control. Comparing the order parameter difference between the two layers, we find that in the experiment, the order parameter values are for both POPE and POPG higher in layer 1 than in layer 2, while the opposite behavior is seen for the control. This indicates that the presence of lipid II leads to more disordered chains of the bilayer lipids POPE and POPG in layer 2. In contrast, its lipid tails cause more ordering in the POPE and POPG lipid tails of layer 1, i.e., the presence of lipid II leads to a more fluid layer 2 and a more viscous, gel-like layer 1. Such altering of the local membrane environment by the presence of lipid II has also been observed in earlier work^46^, but in our case, is connected with the presence of nisin chains. Hence, similar to what is seen for teixobactin^45^, the interactions between nisin and lipid II reduce the integrity of the bilayer.

**Figure 6:**
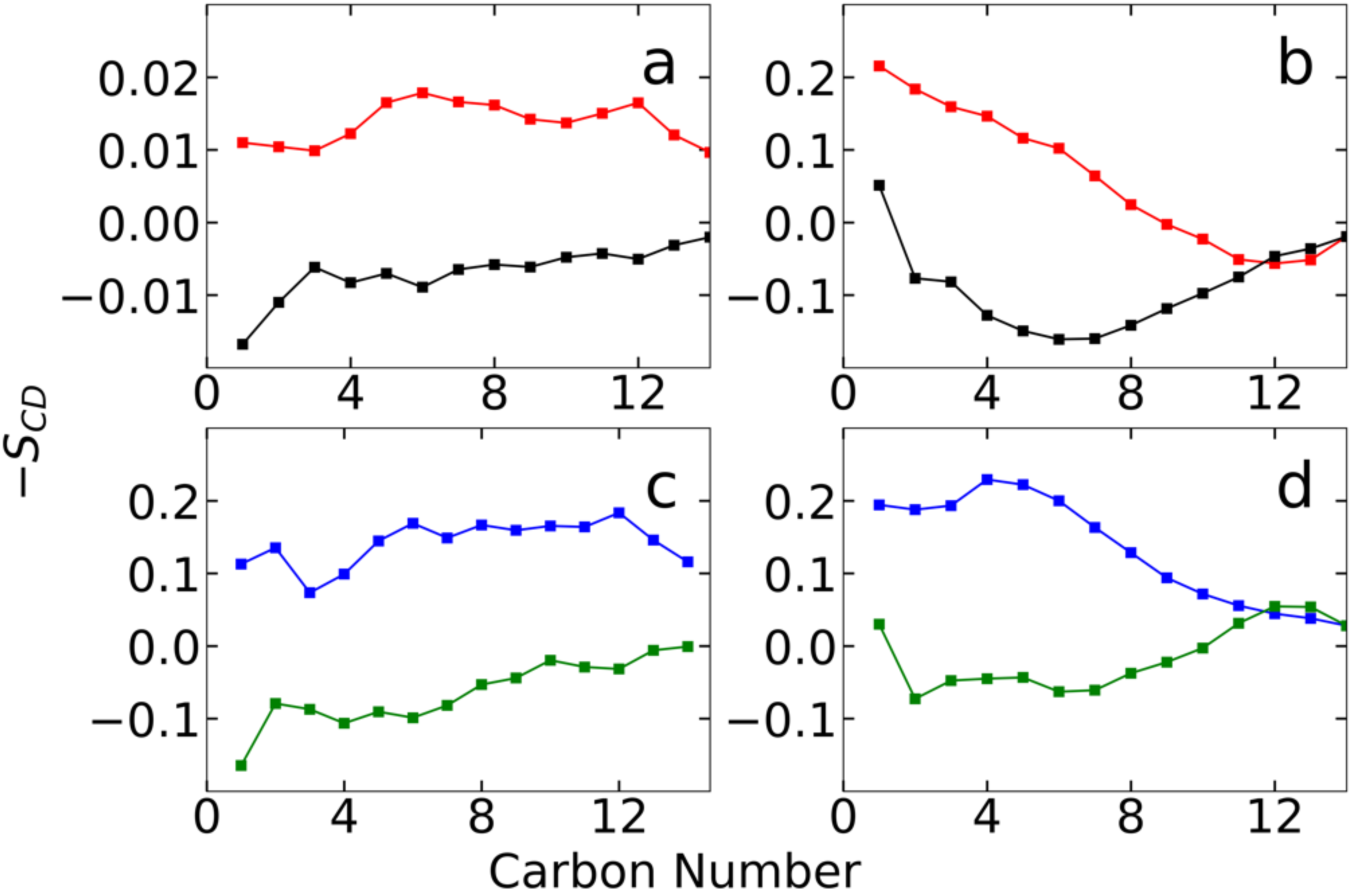
Tail group order parameter -S_CD_, averaged over the final 50 ns of three trajectories and measured separately for POPE and POPG lipids and both layers of the membrane. In (a), we show the differences between the values measured for POPE in the two layers, with the results from simulations in the presence of lipid II drawn in red and that of the control in black. The corresponding plots for POPG are shown in (b). On the other hand, the differences between the order parameter values measured in the experiment and control are shown in (c) for POPE and in (d) for POPG, where we plot in blue the results for layer 1, and in green the results for layer 2.

## III. CONCLUSIONS

Although nisin is non-toxic to humans and has broad and strong antimicrobial activity, its poor solubility and stability limit its use as an antibiotic. Considerable efforts have been put into deriving nisin analogs that overcome these limitations, but equally crucial for the development of future antibiotics is obtaining an in-depth understanding of the mechanisms for nisin’s antimicrobial effectiveness. In order to gain such an understanding, we have conducted large-scale molecular dynamics simulations of a nisin assembly, designed to be consistent with experimental data, that was inserted in a membrane model for gram-positive bacterial cells. As a control, we used the same model of a nisin pore in a bilayer but removed the lipid II molecules. The purpose of our simulations was to study the effect of the bacteria-specific lipid II on the nisin pore stability, and to probe how in the absence and presence of lipid II nisin chains change the bilayer in our model of a bacterial membrane.

We find that contacts formed between the lipid II head group atoms and nisin residues stabilize the geometry of the nisin chains and, as a consequence, the integrity of the pore assembly. Additional stabilization is provided by the proximity of the flexible and moving lipid II tails that constrain the movement of the nisin molecules within the membrane. The pore decays in both absence and presence of lipid II, but more so in the control where lipid II is not present. It should be noted that the lifetime of a pore formed by nisin chains will likely be larger in a bacterial cell than seen in our model which did not account for a flow of water molecules through the pore that would keep the pore open.

The assembly of nisin chains interacting with lipid II chains also changes the property and behavior of the membrane, leading in the neighborhood of the nisin chains to more disordered lipid chains in layer 2. In contrast, it causes more ordering in the POPE and POPG lipid tails of layer 1, where the lipid II chains reside. Of critical importance for these processes are residues 8-11 and 21-28, i.e., rings B and D of the nisin chains. The presence of lipid II leads in both layers locally to an increase in the Area per Lipid (APL), i.e., the density of lipids in the bilayer, with the difference between the two layers much more prominent than in the control. The effect of the increased density is twofold: a local thinning of the bilayer and a spread in the viscosity of the two layers, with layer 2 becoming more fluid and layer 1 (where the lipid II are located) more gel-like. Both effects reduce the stability of the bacterial membrane.

In conclusion, our computational study shows that interaction with lipid II enhances the stability of pores formed by nisin chains in the membranes of gram-positive bacteria, thus increasing the chance for cell leakage. Interactions between nisin chains and lipid II molecules also cause local changes in membrane viscosity and thickness that further destabilize the bacterial membrane. We propose that it is the combination of the two effects that makes nisin an effective antimicrobial agent.

## Supporting information

ZIP file containing start and final configurations, and parameter files for nisin and lipid II

## SUPPLEMENTARY MATERIAL

The Supplementary Material is available free of charge at the URL

A compressed ZIP file contains a README file (.txt), folders with the start configurations (as .gro files) of all trajectories (START), the corresponding final configurations (FINAL), and two .itp files with force field parameters and topologies of the nisin-specific uncommon amino acids and lipid II molecules.

## ACKNOWLEDGMENT

The simulations in this work were done using the SCHOONER cluster of the University of Oklahoma and on TACC resources allocated under grant MCB20016 (National Science Foundation). We acknowledge financial support from the National Institutes of Health under grant GM120578.

## AUTHOR DECLARATIONS

Conflict of Interests

The authors have no conflicts to declare.

## AUTHOR CONTRIBUTIONS

**Miranda S. Sheridan:** Formal analysis (equal); Investigation (lead); Visualization (lead); Conceptualization (equal) Writing – original draft (equal); Writing – review and editing (equal);

**Preeti Pandey:** Conceptualization (equal) Writing – review and editing (equal)

**Ulrich H.E. Hansmann:** Conceptualization (lead); Funding acquisition (lead); Resources (lead); Supervision (lead); Writing – original draft (equal); Writing – review and editing (equal).

## DATA AVAILABILITY

The data supporting this study’s findings are available from the corresponding author upon request for non-commercial use, under the condition that the use of these data is acknowledged.

