## Supplementary material for "In bacterial membranes lipid II changes the stability of pores formed by the antimicrobial peptide nisin": ZIP file containing start and final configurations, and parameter files for nisin and lipid II: README.txt

Directory of files:
# START: This is a folder with the start configurations (as .gro files) ) 
For three trajectories  of the nisin-pore-membrane complex in presence of lipid II (experiment_r1_t0.gro, experiment_r1_t0.gro, experiment_r1_t0.gro), 
and in absence of lipid ii (control_r1_t0.gro, control_r2_t0.gro, control_r3_t0.gro).

#FINAL: This is a folder with the final configurations (as .gro files), taken after 200 ns  
of simulation, for the nisin-pore-membrane complex in presence of lipid II (experiment_r1_t200.gro, experiment_r1_t200.gro, experiment_r1_t200.gro),
and in absence of lipid ii (20.gro, control_r2_t200.gro, control_r3_t200.gro).

# lipa.itp: This file contains the parameters and topology for the lipid II molecules

# nisin.itp:This file contains the parameters and topology describing the uncommon residues in nisin. 

# README.txt: This file
